# Label-Free Proteomics of α-Synuclein-Expressing *Caenorhabditis elegans* Model Uncovers Conserved Pathophysiological Pathways in Parkinson’s Disease

**DOI:** 10.64898/2026.09.17.752360

**Authors:** Iverson Conrado Bezerra, Maria Luiza de Lima Vitorino, Josivan Barbosa de Farias, Maria Gabriela Arruda de Medeiros, Roberto Afonso da Silva, José Luiz de Lima Filho, Gustavo José da Silva Pereira, Priscila Gubert

## Abstract

Parkinson’s disease (PD) is the second most prevalent neurodegenerative disease in the elderly population, characterized by a wide variety of motor and non-motor symptoms. Its complex biochemical and molecular mechanisms make it challenging to study PD directly in patients or to replicate it in vitro, underscoring the need for careful model characterization to advance translational research. The free-living nematode *Caenorhabditis elegans* has been widely used in neuroscience studies. Technological advancements enabled the development of transgenic strains that mimic aspects of neurodegenerative diseases, such as the OW13 strain [*grk-1*(ok1239) X; *pkIs2386* IV (*unc-54*p::alpha-synuclein::YFP + *unc-119*(+))], which expresses human α-synuclein in body wall muscle cells in a sensitized G-protein coupled receptor kinase-deficient background. This study conducted an exploratory proteomic analysis of the OW13 strain. A cross-species comparative analysis using post-mortem substantia nigra proteomics from PD individuals identified a convergent signature of 76 conserved *C. elegans* proteins, corresponding to 70 human orthologs, derived from the 373 DAPs. Among these conserved candidates, 68 proteins were upregulated, and 8 were downregulated. Functional enrichment linked these DAPs to the tricarboxylic acid cycle, mitochondrial function, and protein quality control, suggesting that α-synuclein aggregation triggers a broad proteostatic and bioenergetic remodeling resembling PD pathophysiology. This first proteomic and cross-species characterization of the OW13 strain strongly supports *C. elegans* as a robust PD model.

## 1. Introduction

Population aging worldwide is a major challenge, as it increases the incidence and prevalence of various age-related diseases.^1^ Among the variety of conditions in the elderly population, Parkinson’s disease (PD) is the second most common progressive neurodegenerative disease.^2^ The occurrence of PD has doubled in the last 25 years,^3^ with projections of 25.2 million individuals by 2050, representing a significant 112% increase, and an estimated 216 cases per 100,000 inhabitants.^4^ The economic burden associated with PD management has reached US$52 billion per year in the USA, with projected increases due to its progressive nature.^5^

PD pathophysiology is characterized by the intracellular accumulation of α-synuclein, a hallmark supported by SNCA mutations as the first genetic cause of familial PD in 1997,^6^ leading to progressive degeneration of dopaminergic neurons in the substantia nigra pars compacta.^2^ Consequently, PD patients exhibit several locomotor symptoms, such as tremor, rigidity, bradykinesia, and postural instability, as well as some non-motor signs and symptoms.^7^ The motor circuit is compromised by α-synuclein accumulation in dopaminergic neurons,^8^ as well as having a potential role in regulating peripheral nerves and the function of the skeletal muscles innervated by them.^9^

α-synuclein is a highly dynamic protein that can adopt multiple conformational states, ranging from physiological monomers and tetramers to pathological oligomers, protofibrils, and fibrillar aggregates. The biological consequences of α-synuclein accumulation are strongly influenced by its structural conformation, as distinct conformers exhibit different aggregation propensities, intracellular distributions, and affinities for cellular proteins.^10^ As a result, specific α-synuclein species establish unique interaction networks that differentially affect processes such as synaptic transmission,^11^ protein homeostasis,^12^ mitochondrial function,^13^ and cellular signaling. The subcellular localization of these conformers further shapes their pathogenic potential, exposing them to distinct molecular partners and contributing to the heterogeneous cellular responses observed in PD.

Under pathological conditions, several post-translational modifications of α-synuclein, including phosphorylation, ubiquitination, and truncation, promote the conversion of physiological conformers into aggregation-prone species, favoring oligomer formation and the accumulation of insoluble fibrils in neurons.^14^ These pathogenic assemblies subsequently impair multiple cellular processes through aberrant protein interactions and organelle dysfunction, particularly affecting mitochondrial homeostasis and redox balance, which contribute to oxidative stress and neurodegeneration in PD.^15, 14^

Experimental models play an essential role in elucidating PD pathophysiological mechanisms and supporting preclinical research aimed at dissecting disease-associated molecular pathways.^16^ Several mammalian species, including rats, mice, and non-human primates, have been widely used to investigate PD-related mechanisms and cellular alterations.^17,16^ In addition, invertebrate organisms such as *Caenorhabditis elegans* (*C. elegans*) (Maupas, 1900) have also been employed as experimental models.^18^ Although their physiology differs considerably from that of humans, these organisms provide valuable insights into conserved molecular and cellular mechanisms associated with neurodegeneration.^19^ Nevertheless, the translational relevance of experimental models in PD remains debated, as currently available models do not fully reproduce the complex and progressive pathology observed in humans, which may limit the direct extrapolation of preclinical findings to clinical settings.^20^ Therefore, a deeper characterization of disease models, together with a clearer understanding of their strengths and limitations, remains essential for improving translational research in PD.^21^

*C. elegans* is a free-living nematode extensively employed as a model organism in studies of cell and developmental biology.^18^ Moreover, *C. elegans* shows substantial genetic conservation with humans, with ∼38% of its genes having human orthologs and ∼83% proteome homology.^22–24^ Advances in biotechnology have expanded its use through transgenic approaches, enabling the generation of strains that model human diseases, such as PD.

The transgenic *C. elegans* OW13 model harbors the integrated transgene *pkIs2386* [*unc-54*p::alpha-synuclein::YFP + *unc-119*(+)] in a *grk-1*(ok1239) loss-of-function background and expresses human α-synuclein fused to yellow fluorescent protein in body wall muscle cells.^25^ The expression of α-synuclein in muscle cells facilitates subcellular visualization, owing to the relatively large size of muscle cells.^25^ Furthermore, the model exhibits α-synuclein inclusions, recapitulating the formation of insoluble protein aggregates characteristic of PD pathology.^25^ In *C. elegans*, α-synuclein deposits generate proteotoxicity,^26^ reduced lipid content,^27,28^ increased oxidative stress,^28^ reduced chemotaxis,^29^ and cause mitochondrial dysfunction,^28^ some relevant findings in the pathophysiology of PD.

In the present study, we performed an exploratory proteomic analysis to profile the proteome of a *C. elegans* PD model (OW13) expressing human α-synuclein in muscle cells. In parallel, we analyzed proteomic data from the post-mortem substantia nigra of individuals with PD. We conducted a cross-species comparative analysis to map orthologous alterations and identify shared, conserved α-synuclein-associated pathways across species, providing a molecular baseline for understanding systemic proteostatic and bioenergetic stress.

## 2. Material and Methods

### 2.1. Proteomics Analysis

#### 2.1.1. C. elegans Strain and Maintenance

The wild-type N2 strain and PD model OW13 [*grk-1*(ok1239) X; *pkIs2386* IV *(unc-54*p::alpha-synuclein::YFP + *unc-119*(+))] of *C. elegans* were obtained from the *Caenorhabditis* Genetics Center (CGC, Minnesota, USA). OW13 animals express human α-synuclein in the body wall muscles. The YFP tag is fused to the C-terminus of α-synuclein. Worms were maintained and cultivated on Petri dishes containing nematode growth medium (NGM) composed of 3 g NaCl, 2.5 g peptone, 17 g agar, and 975 mL autoclaved H_2_O; supplemented with 1 mL of 1 mol L^−1^ CaCl_2_, 1 mL of cholesterol solution (5 mg mL^−1^ in ethanol), 1 mL of 1 mol L^−1^ MgSO_4_, and 25 mL of 1 mol L^−1^ KPO_4_ buffer. Animals were fed *Escherichia coli* (*E. coli*) OP50 and maintained at a controlled temperature of 20 °C, with humidity above 95%. We synchronized the population to ensure all worms were at the same larval stage, and treated pregnant hermaphrodites with a bleaching solution (1 M NaOH, 1% NaClO, and distilled H2O) to disrupt the cuticle and facilitate egg release. Eggs were then maintained in M9 buffer (3 g L^−1^ KH_2_PO_4_, 6 g L^−1^ Na_2_HPO_4_, g L^−1^ NaCl, and 1 mM MgSO_4_) until hatching and development to the L1 larval stage at 20 °C.

#### 2.1.2. Validation of Transgenic Animals by Fluorescence Microscopy

Transgenic animals were validated by fluorescence microscopy prior to subsequent experimental procedures. The *C. elegans* PD model (OW13) at the L4 larval stage was transferred from NGM culture plates onto glass microscope slides containing a 2% agarose pad. The animals were immobilized with 1 M sodium azide and covered with a coverslip. Fluorescence images were acquired using a Zeiss Axio Imager (Carl Zeiss AG, Oberkochen, Germany). Animals were examined for the presence of fluorescent α-synuclein inclusions in body wall muscle cells, confirming transgene expression (Fig. S1).^28^

#### 2.1.3. Protein Sample Processing and Digestion

Approximately 15,000 L4 worms were used per biological replicate, with three independent biological replicates per group.^30^ Samples were mechanically disrupted by manual maceration using the Sample Grinding KitTM (Cytiva, Marlborough, MA, USA) according to the manufacturer’s protocol and using a Lysis buffer containing 7 M urea, 2 M thiourea, 4% 3-[(3-cholamidopropyl)dimethylammonio]-1-propanesulfonate (CHAPS), 40 mM DTT, and 0.5% Immobilized pH Gradient Buffer. During extraction, a protease inhibitor cocktail (Promega, Madison, WI, USA) was added at a protein ratio of 1:50 (w/w). The lysates were then centrifuged at 10,000 × g for 5 min, and the resulting supernatant was collected.

Protein concentration was subsequently determined using the 2-D Quant Kit™ (Cytiva, Marlborough, MA, USA). 100 μg of total protein from each group was subsequently processed using the 2-D Clean-Up™ Kit protocol (Cytiva, Marlborough, MA, USA) to remove interfering contaminants. After purification, 100 μg of protein from each group was resuspended in 8 M urea. Proteins were reduced with 100 mM dithiothreitol at 30°C for 30 min, then alkylated with 300 mM iodoacetamide at 30 °C for 30 min in the absence of light. The solution was then diluted by adding 50 mM NH HCO. The digestion was performed using sequencing-grade modified trypsin (Promega, Madison, WI, USA) at a protein/protein ratio of 1:50 (w/w), followed by incubation at 37 °C for 18 h. After digestion, tryptic peptides were centrifuged at 11,000 × g for 10 min at 4 °C. The supernatant was concentrated at 30 °C using a SpeedVac concentrator 5301 (Eppendorf, Hamburg, Germany).^31^

#### 2.1.4. nanoUltraperformance Liquid Chromatography-mass Spectrometry/mass Spectrometry (nUPLC-MS/MS) Analysis

The tryptic peptides were recovered in 0.1% formic acid to obtain a final protein concentration of 1 μg μL^−1^ and transferred to total recovery vials. Peptide separation was performed using an M-Class ACQUITY UPLC system (Waters Corporation, Milford, MA, USA). The system was configured with a single-pump trap column (nanoEase M/Z Symmetry C18, 100 Å, 5 μm, 180 μm × 20 mm) operating at a flow rate of 0.3 μL/min□^1^ and coupled to an analytical column (nanoEase M/Z HSS C18 T3, 100 Å, 1.8 μm, 75 μm × 250 mm) operating at the same flow rate and maintained at 40 °C. The mobile phase A consisted of water containing 0.1% (v/v) formic acid, while mobile phase B consisted of acetonitrile containing 0.1% (v/v) formic acid. Peptide elution was carried out over 133 min, with a loading time of 10 min, using the following gradient: (I) 0–8 min, 95% A and 5% B; (II) 8–98 min, 60% A and 40% B; (III) 98–103 min, 15% A and 85% B; (IV) 103–108 min, 15% A and 85% B; (V) 108–109 min, 95% A and 5% B; and (VI) 109–133 min, 95% A and 5% B (initial conditions).

The UPLC system was coupled to a Synapt XS quadrupole Q-ToF mass spectrometer with T-Wave ion mobility (Waters Corporation, Wilmslow, UK), operating at a mass resolution of 30,000 full width at half maximum. The electrospray low-flow probe was operated at a capillary voltage of 3 kV, a sampling cone voltage of 40 V, and a source offset of 30 V. The source temperature was set to 100 °C, and the cone gas flow was 50 L min^−1^. The time-of-flight mass analyzer was externally calibrated using a NaCsI mixture over m/z 50 to 2000. A lock-mass reference signal of Glu-fibrinopeptide B (m/z 785.8426) was acquired every 30 s. Data were acquired using Ultra-Definition Mass Spectrometry in positive ion mode, with each biological replicate analyzed in technical duplicate.^31^

#### 2.1.5. Data Processing, Protein Identification, and Quantification

Protein identification was carried out from nUPLC–MS/MS data using Progenesis QI software version 4.7. Database searches were performed against the *C. elegans* proteome obtained from UniProt (downloaded on 16/09/2025),^32^ including both reviewed and unreviewed entries. The parameters applied to spectrum processing and database searching included allowance for missed trypsin cleavages, a maximum protein mass of 750 kDa, carbamidomethylation of cysteine as a fixed modification, and methionine oxidation as a variable modification. The search criteria required at least 2 fragment-ion matches per peptide, 5 fragment-ion matches per protein, and 2 unique peptides per protein. Protein identifications were filtered using a false discovery rate of 1%. Relative quantification was performed using the Hi3 label-free method, based on the three most abundant peptides for each identified protein. The algorithm considered only proteins with frequency scores and confidence intervals exceeding 99% as valid identifications.^31^

#### 2.1.6. Bioinformatics Analysis of Proteomics Results

Progenesis QI software version 4.7 was used for protein identification, normalization, and protein group extraction. DAPs were assessed using analysis of variance (ANOVA) with a significance threshold of *p* < 0.05. Additional analyses were performed using the Omicscope algorithm version 1.4.2 and R statistical software.^33^ Principal component analysis (PCA) was performed to assess sample clustering, and volcano plots were generated to distinguish and visualize up- and downregulated proteins. The Fold change (FC) values were log_2_-transformed, and adjusted *p*-values were converted to -log^10^. Volcano plots were generated using log_2_ FC cutoffs of ≤ −1.2 and ≥ 1.2 with a significance threshold of *p-adjusted* < 0.05. The average coefficient of variation (CV) was used to assess reproducibility among replicates, calculated from normalized abundance.^34, 35^ The CV was calculated according to the following equation (1):

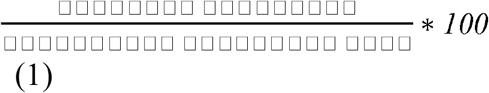

Figure 1H provides a simplified schematic of the methodological process.

**Figure 1.**
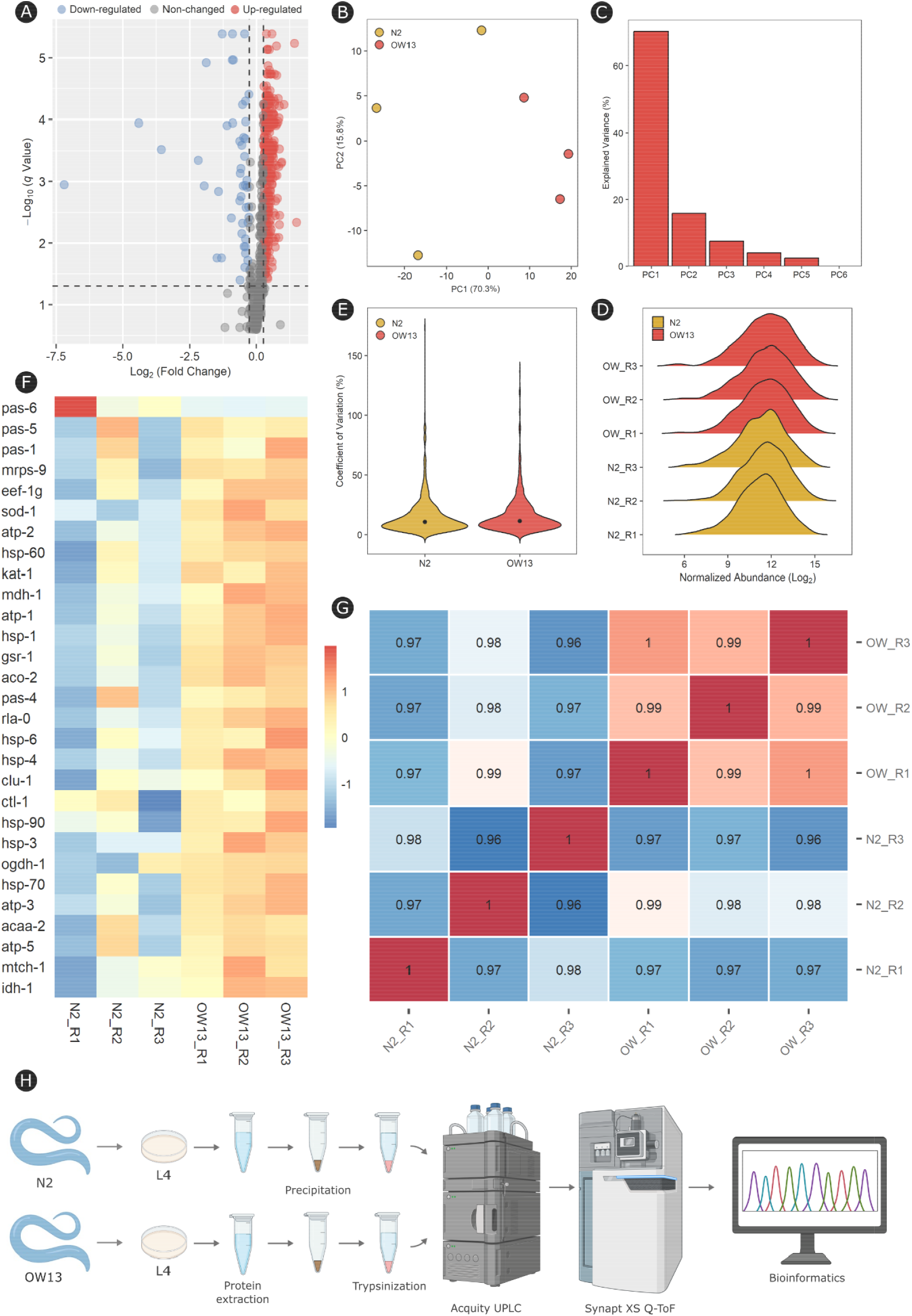
Proteomic analysis of *C. elegans* strain OW13. (A) Volcano plot of the proteomic profile of *C. elegans* PD animal (OW13) compared to wild-type, with Log_2_ FC of ≤ −1.2 and ≥ 1.2 and a significance threshold of *p-adjusted* < 0.05. (B) Principal component analysis. (C) Percentage of variance explained by principal components. (D) Data normalization. (E) Coefficient of variation across replicates. (F) Heatmap highlighting DAPs with repercussions on PD. (G) The correlation coefficient between groups and replicates. (H) Simplified schematic representation of protein extraction, digestion, identification, and analysis.

### 2.2. Cross-Species Comparative Analysis

#### 2.2.1. Parkinson’s Disease-Associated Differentially Expressed Proteins and Source Datasets

The proteome of individuals with PD used was obtained from *Jang Y et al.,* who kindly provided the dataset.^36^ Quantitative proteomic analysis was performed on post-mortem tissue from the substantia nigra, a brain region critically affected by the progressive degeneration of dopaminergic neurons. Brain tissue samples were obtained from the *Brain Resource Center* at *Johns Hopkins University School of Medicine*. The diagnosis of PD was established according to the *UK Brain Bank* clinical criteria and subsequently confirmed by neuropathological analysis. Control subjects showed no clinical or neuropathological evidence of neurodegenerative diseases. The original publication provides detailed clinical information on patients and control subjects.^36^

In the original study, the proteomic profile was obtained using quantitative proteomics by high-resolution mass spectrometry, enabling large-scale identification and quantification of proteins in substantia nigra samples. In the present study, the processed protein dataset was used.^36^ DAPs were filtered, and only those with *q* values *< 0.05* were included in our analyses and used for comparative analyses and cross-species integration with the proteomic dataset from the *C. elegans* PD model.

#### 2.2.2. Cross-Species Identification of Human Orthologs of C. elegans Proteins

The study workflow was adapted from *Ray et al.*.^37^ The *C. elegans* DAPs were filtered according to log_2_ FC ≥ 1.2 and ≤ −1.2 and significance thresholds (adjusted *p*-value < 0.05). *C. elegans* protein symbols were submitted to OrthoList 2, a database containing a curated compilation of *C. elegans*-human orthologs obtained through meta-analysis.^38^ The human orthologs were compared with the DAPs identified in the PD dataset.^36^ Human proteins present in both datasets were identified, and their corresponding *C. elegans* proteins were retrieved, yielding a final dataset of conserved proteins used for subsequent analyses.

Comparative analyses were performed in Python 3.10 to identify overlapping proteins between human orthologs derived from *C. elegans* proteomics and regulated proteins in PD.^36^ Proteins present in both datasets were considered conserved candidates and were used in subsequent analyses.

#### 2.2.3. Gene Ontology and Cross-species pathway conservation analysis

Functional enrichment analyses were performed using the Gene Ontology (GO) database, considering the categories Biological Process (BP), Cellular Component (CC), and Molecular Function (MF), as well as the Kyoto Encyclopedia of Genes and Genomes (KEGG) pathway database. Human proteins identified in PD individuals and their orthologous proteins identified in *C. elegans* through cross-species analysis were subjected to functional enrichment using the DAVID platform.^39^ This analysis was conducted to investigate the functional enrichment and conservation of molecular pathways between the *C. elegans* PD model and individuals with PD.^37^

#### 2.2.4. Parkinson’s Disease PPI Network and Core Proteins

The PPI network was constructed to investigate the potential regulatory roles of the human proteins. Thus, only human proteins conserved with the *C. elegans* proteomic results were included in the network construction. Interaction data were obtained from the STRING database.^40^ The STRING interaction network was imported into Cytoscape version 3.10.4, applying a confidence score cutoff of 0.4.^41^ Isolated nodes were removed, and the resulting network was used for downstream analysis. The CytoHubba plugin was used to evaluate the network’s topological properties.^42^

Nodes with high connectivity and centrality in the PPI network were considered hub proteins. To identify key regulators, Degree, Closeness Centrality (C_C_), and Betweenness Centrality (C_B_) were calculated. Subsequently, a Venn diagram was generated to identify the common hub proteins shared among the top-ranked nodes identified by Degree, C_C_, and C_B_.

#### 2.2.5. Ortholog-Based PPI Network Construction in C. elegans

Animal models of human diseases are widely used to investigate conserved molecular mechanisms underlying pathological processes. The identification of orthologous proteins between humans and *C. elegans* may reveal conserved biological functions and regulatory pathways associated with PD.^43^ Therefore, a PPI network was constructed using the subset of *C. elegans* DAPs that presented human PD orthologs. The interaction network was generated using the STRING database (https://string-db.org/) and visualized in Cytoscape version 3.10.4 using the Omics Visualizer plugin, and applying a confidence score cutoff of 0.4.^40,44^ After removing isolated nodes, the resulting network was used for downstream analysis.

Topological properties of the network were evaluated using the CytoHubba plugin.^42^ The Degree, C_C_, and C_B_ measurements were calculated to identify the most influential proteins within the interaction network. Subsequently, a Venn diagram was generated to identify overlapping hub proteins among the top-ranked nodes identified by Degree, C_C_, and C_B_. These overlapping proteins were identified as central candidates and used in subsequent analyses.

#### 2.2.6. Identification of key regulators in the C. elegans orthologous network

After identifying PD and *C. elegans* proteins based on Degree, C_C_, and C_B_, a search was conducted to identify key regulators shared between the organisms and their respective interaction networks.^37^ Proteins with high connectivity and centrality values were considered hub candidates and evaluated for their potential regulatory roles within the PPI networks. These hubs, which contribute to the structural organization and functional backbone of interaction networks, were further examined as important regulators of PPI interactions.

#### 2.2.7. Analysis of modules in the C. elegans orthologous network

Network modules correspond to groups of proteins that interact more frequently among themselves than with proteins outside the group.^45^ To identify these clusters within the PPI network, the MCODE (Molecular Complex Detection) plugin, version 2.0.3, was used to detect densely connected regions that may represent functional protein complexes.^46^ The parameters used for the MCODE analysis were: Degree cut-off = 2, node score cut-off = 0.2, k-score = 2, and max depth = 100. Only modules with a score ≥ 2 were retained and considered significant for further analysis.^37^

## 3. Results and Discussion

The incompletely understood etiology of PD underscores the need for well-characterized experimental models to elucidate the conserved cellular and molecular cascades triggered by α-synuclein pathology. Although the disease involves proteins, these models have particularly contributed to understanding pathological features such as α-synuclein aggregation, mitochondrial dysfunction, proteostasis imbalance, and neuronal degeneration.

To capture these pathological cascades, *C. elegans* serves as a well-established model system that exhibits substantial conservation with human molecular pathways associated with PD pathophysiology, particularly in proteostatic regulation and mitochondrial bioenergetics. Specifically, the transgenic *C. elegans* PD model (OW13) is employed to screen for molecules with anti-aggregant potential, allowing evaluation of α-synuclein-induced content and toxicity,^47^ stress resistance,^48^ and locomotor behavior.^48^ As originally established by van Ham *et al*., the OW13 strain combines muscle-directed expression of human α-synuclein with a loss-of-function of *grk-1*, which acts as a specific genetic modifier that decreases the accumulation of α-synuclein inclusions. Consequently, by comparing OW13 against the N2 strain, our label-free quantitative proteomic profiling captures the integrated cellular and proteostatic response to human α-synuclein expression within a sensitized, GRK-1-deficient molecular context.^25^

We performed proteomic profiling at the L4 larval stage to characterize early molecular alterations induced by α-synuclein before the onset of generalized organismal decline. Because physiological aging independently alters the nematode proteome, profiling at later adult stages could confound pathology-associated molecular signatures with age-related changes.^49,50^ Therefore, we selected the L4 stage to minimize age-associated proteomic interference while capturing active molecular and compensatory stress responses associated with α-synuclein expression. We subsequently applied proteomics coupled with cross-species bioinformatics to determine whether this genetically defined model recapitulates core pathological pathways observed in post-mortem PD substantia nigra tissue.

Firstly, following the extraction and identification of the proteins, the DAPs of the *C. elegans* PD model (OW13) were identified using a log_2_ FC cutoff of ≥ 1.2 and ≤ −1.2 (*p-adjusted < 0.05*). Quantitative analysis identified 497 proteins with significant abundance changes. Applying the established log_2_ FC threshold, a final set of 373 DAPs was defined, with 326 proteins significantly upregulated and 47 significantly downregulated (Fig. 1A; Table S1).

Rigorous quality-control analyses ensured that the observed proteomic variations reflected true biological responses rather than technical artifacts. The PCA showed a distinction between control animals and the transgenic PD model, with PC1 accounting for 70.3% and PC2 for 15.8% of the explained variance (Fig. 1B-C). Data normalization was applied to enable more accurate comparisons, presenting mean abundances with corresponding error ranges across all conditions and replicates (Fig. 1D). High data reproducibility was observed in the analyses, along with a wide range of identified and quantified proteins, as evidenced by the dynamic range (Fig. S2).

Furthermore, the robustness of the overall analytical workflow was supported by a low mean CV for detected proteins, at 22% in control animals and 10% in the PD model (Fig. 1E). The CV observed across replicates remained low, indicating high reproducibility of the measurements, consistent with values commonly reported in previous proteomic analyses of *C. elegans*.^30, 34, 35^ In contrast, the higher variability observed for a protein fraction may be safely attributable to differences in sampling times across runs and/or limitations in MS/MS detection of low-abundance proteins. To visually explore these regulated protein patterns across replicates, a heatmap was generated, providing a structured overview of the proteins essential to addressing the fundamental hypotheses of our study (Fig. 1F).

Pearson correlation analysis of replicate samples further corroborated internal reliability, showing strong concordance among biological replicates and ruling out potential technical variability or normalization artifacts (Fig. 1G). Therefore, high correlation coefficients were observed among replicates from the same experimental group, indicating robust reproducibility and validating the dataset for deeper analysis.

After establishing the statistical robustness of the data, we proceeded to analyze the molecules’ physical properties. The molecular mass of the identified proteins showed peaks at ∼20 kDa and ∼50 kDa, although the proteins extend up to ∼110 kDa (Fig. 2A). Furthermore, the findings regarding the mass of the identified proteins corroborate the amino acid (aa) sequence length, with the vast majority of proteins smaller than 500 aa, peaking near 200 aa, indicating the predominance of medium-sized proteins in the dataset (Fig. 2B). The peptide identification score revealed density peaks at approximately 0.5 × 10^1^, suggesting consistency in the peptide population (Fig. 2C). The retention time distribution showed a higher detection concentration between 0.3 × 10^1^ and 0.6 × 10^1^, a behavior consistent with chromatographic separation patterns (Fig. 2D). Finally, correlations were identified between molecular weight and the number of unique peptides (Fig. 2E), peptide count (Fig. 2F), confidence score (Fig. 2G), and sequence coverage (Fig. 2H).

**Figure 2.**
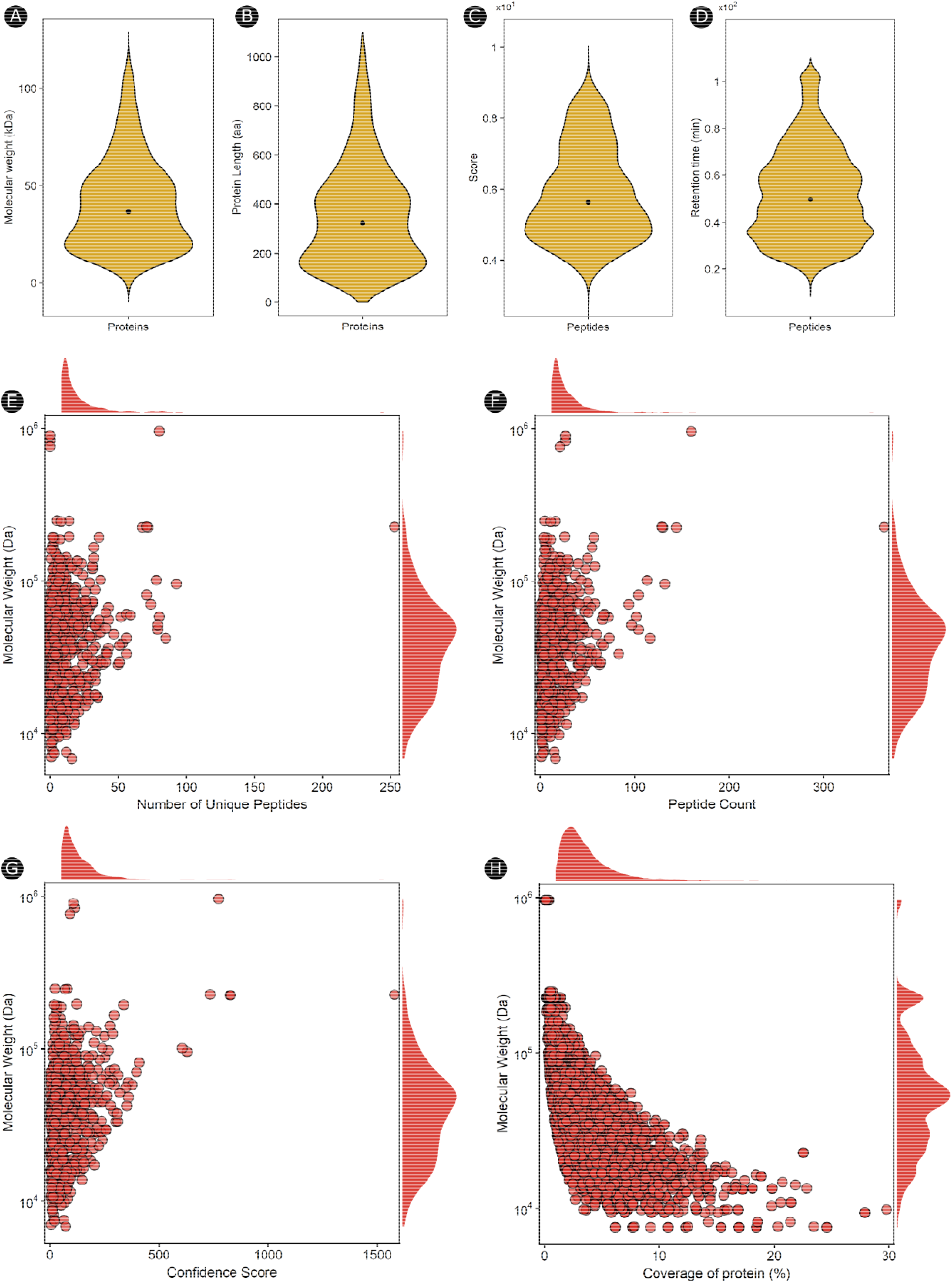
Quality control and physicochemical characteristics of the total identified proteome. Global distributions of peptide and protein metrics across all combined samples to evaluate technical nUPLC-MS/MS, nano-ultraperformance liquid chromatography-mass spectrometry/mass spectrometry performance rather than biological contrast. (A) Protein molecular mass (kDa). (B) Protein sequence length (amino acids). (C) Fragmentation identification score. (D) Peptide retention time (min). (E) Molecular weight versus number of unique peptides. (F) Molecular weight versus peptide count. (G) Molecular weight versus confidence score. (H) Sequence coverage (%) relative to target protein molecular weight.

To establish a translational framework linking the *C. elegans* PD model to human disease, we integrated our findings with PD datasets. In post-mortem substantia nigra tissue from individuals with PD, *Jang* and coworkers previously detected proteins consistent with the molecular characteristics of PD.^36^ In the PD dataset, 1,383 proteins with a *q-value <* 0.05 were used in the cross-species analysis to identify orthologs of human PD-related proteins in *C. elegans*. Given that comparative genomics has shown that ∼83% of the *C. elegans* proteome has homologous genes in humans,^37^ we systematically mapped our nematode DAPs filtered by log_2_ FC (373 DAPs) using OrthoList 2, which successfully yielded 650 human orthologs (Table S2).

Cross-referencing of 650 orthologs with the proteomic dataset from PD individuals identified 76 *C. elegans* proteins corresponding directly to 70 human orthologs,^36^ revealing distinct one-to-many (1:*m*) orthology relationships. This cross-species overlap was statistically significant under a hypergeometric test (Fisher’s exact test, *p* = 0.00015, Odds Ratio = 1.7), assuming a reference human proteome of 20,000 genes.^51^ Based on our established log_2_ FC cutoffs, 68 of these 76 *C. elegans* proteins were upregulated, while 8 were downregulated (Table S3). Collectively, the convergence of these independent datasets highlights conserved cross-species molecular candidates that may serve as prioritized targets for future functional and mechanistic investigation in PD.

To investigate the function of the 76 conserved proteins from the initial 373 DAPs dataset, we performed a functional enrichment analysis using DAVID.^39^ In BP, the proteins were significantly annotated for protein folding (*p* < 0.0001), protein refolding (*p* < 0.001), and tricarboxylic acid cycle (*p* < 0.001) (Fig. 3A-B, Table S4). CC revealed that the enriched terms were predominantly associated with mitochondria (*p* < 0.001), ribosomes (*p* < 0.0001), and cytoplasm *(p* < 0.0001) (Fig. 3A-B, Table S4). MF exhibited significant enrichment for heat shock protein binding (*p* < 0.001), protein folding chaperone (*p* < 0.001), and ATP-dependent protein folding chaperone (*p* < 0.001) (Fig. 3A-B, Table S4). Furthermore, we evaluated KEGG pathways, and the proteins were then associated with the citrate cycle (TCA cycle; *p* < 0.01), carbon metabolism (*p* < 0.001), motor proteins (*p* < 0.0001), and protein processing in endoplasmic reticulum (*p* < 0.05) (Fig. 3A-B, Table S4).

**Figure 3.**
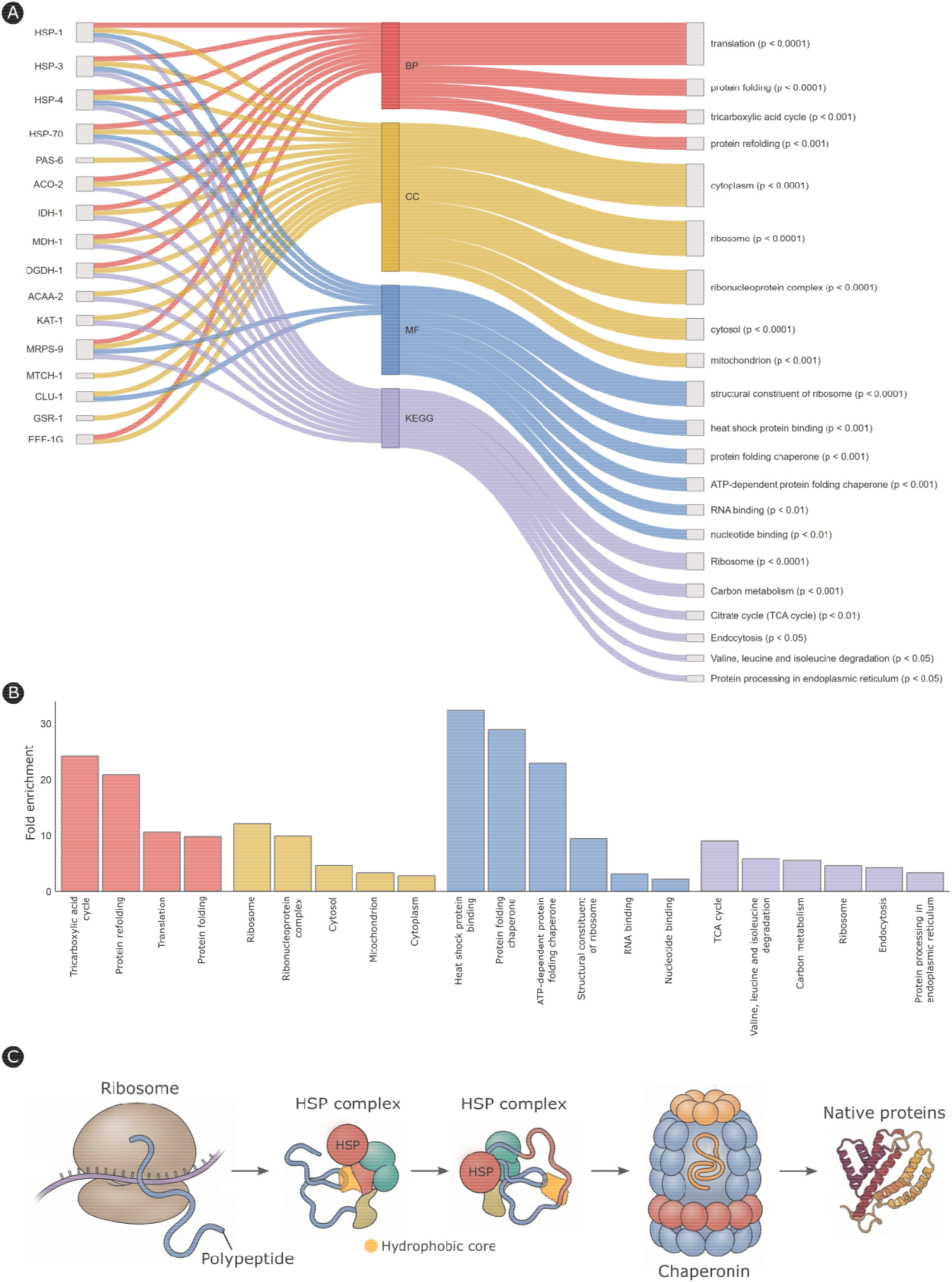
Enrichment analysis of the *C. elegans* PD model (OW13). (A) The diagram illustrates enrichment terms for proteins in biological processes (BP), cellular components (CC), molecular functions (MF), and metabolic pathways (KEGG). (B) Bar chart displaying fold enrichment, with each colored group corresponding to a functional domain. (C) Schematic representation of the protein folding process mediated by molecular chaperones. Ribosomes translate mRNA to synthesize polypeptide chains. Polypeptides interact with HSPs that transiently bind their hydrophobic cores, stabilizing the structure and guiding its conformation. These substrates are then transferred to chaperonin complexes, where they are folded, and a functional protein is released. ^55^ Only the proteins directly discussed in the paper are depicted; the complete analysis is provided in the supplementary tables.

The biological patterns revealed by the functional enrichment analysis indicate that disruption of protein homeostasis and impairment of cellular energy metabolism are the major, interconnected processes associated with PD pathophysiology. The DAPs associated with these pathways provide insight into the molecular mechanisms underlying disease.

The enrichment of chaperone-associated terms across multiple annotation categories is consistent with the well-established involvement of proteostatic dysfunction in PD. Proteostress is a prominent feature of the disease,^52^ which the accumulation of α-synuclein promotes, triggering toxic cellular responses that lead to neuronal dysfunction and degeneration. These alterations are closely associated with disruptions in proteostasis, the cellular network responsible for maintaining protein folding, stability, and degradation^53^. Consequently, molecular processes involved in protein folding, chaperone activity, and protein quality control have increasingly been recognized as central components of PD.^52,54^ Our exploratory analysis, therefore, revealed significant enrichment of *C. elegans* proteins associated with protein homeostasis, including co-chaperones, proteasome, and heat shock proteins (HSPs).

Consistent with the activation of proteostasis-related processes, the HSPs HSP-1 (*p* < 0.0001), HSP-3 (*p* < 0.0001), HSP-4 (*p* < 0.0001), HSP-70 (*p* < 0.0001), HSP-60 (*p* < 0.001), HSP-6 (*p <* 0.01), and HSP-90 (*p < 0.01*) were upregulated in the *C. elegans* PD model (Fig. 4A-G). Reflecting their roles in directing protein folding and refolding, these molecular chaperones were annotated for BP (Fig. 3A; Table S4) and MF (Fig. 3A; Table S4). These proteins were also associated with protein processing in the endoplasmic reticulum pathway in KEGG (Fig. 3A; Table S4), highlighting their involvement in protein quality control mechanisms. In the dataset from PD individuals, the human orthologs HSPA1B and HSPA8/hsc70 (70-kDa heat shock cognate), corresponding to the *C. elegans* HSP-1, 3, 4, and 70, were upregulated. This increase suggests the activation of compensatory proteostasis mechanisms associated with α-synuclein aggregation.

**Figure 4.**
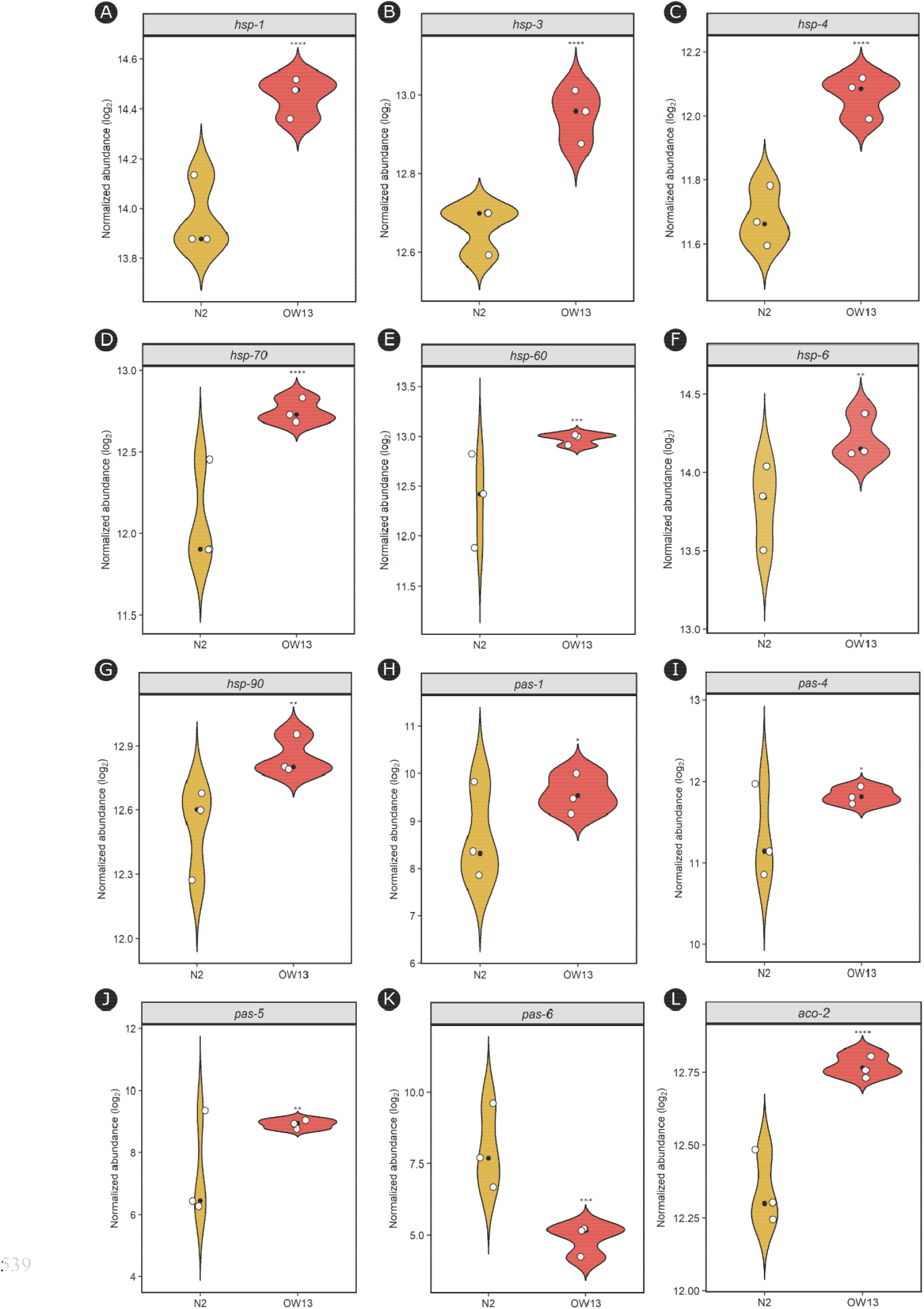
Violin plots of the normalized abundance of DAPs in N2 *wild-type* and *C. elegans* PD model (OW13). (A) Heat shock protein 1. (B) Heat shock protein 3. (C) Heat shock protein 4. (D) Heat shock protein 70. (E) Heat shock protein 60. (F) Heat shock protein 6. (G) Heat shock protein 90. (H) Proteasome subunit alpha type-6. (I) Proteasome subunit alpha type-7. (J) Proteasome subunit alpha type-5. (K) Proteasome subunit alpha type-1. (L) aconitase-2. Data are displayed as distribution plots, with median values. Statistical significance was assessed using an ANOVA. \**p□<*□0.05, \*\**p□<*□0.01, \*\*\**p□<*□0.001, **** *p□<*□0.0001.

Mechanistically, α-synuclein aggregation in PD impairs proteostasis and triggers cellular protein quality control, in which HSPs play a central role. Acting as molecular chaperones, HSPs assist in native protein folding (Fig. 3C), promote the refolding of misfolded proteins, and prevent the accumulation and aggregation of damaged species.^12^ When proper refolding cannot be achieved due to cellular stress, these chaperones facilitate the targeting of damaged proteins to degradation pathways, such as the ubiquitin-proteasome system or autophagy, attempting to maintain proteostasis under pathological conditions (Fig. 3C).^56, 57^

The upregulation of HSPs in the *C. elegans* PD model, alongside the increased abundance of the human orthologs HSPA1B and HSPA8 in our comparative dataset, points to a robust shared compensatory response across species. This increase in HSP family member abundance in PD is consistent with a critical protective response to the accumulation of abnormal proteins, facilitating their degradation and clearance.^58^ In particular, HSP70 family proteins have been shown to attenuate dopaminergic neuronal damage in experimental PD models, including *Drosophila melanogaster* and rats.^59, 60^

Furthermore, recent findings demonstrated that overexpression of HSP70 significantly mitigated α-synuclein-induced neurodegeneration and motor impairment in rodents.^59, 61^ Similarly, HSP60, the ortholog of HSP-60 in *C. elegans*, directly regulates oxidative stress and free radicals, both of which occur mainly in mitochondria.^62^ Thus, the evolutionary conservation of these HSPs between *C. elegans* and PD individuals reinforces the translational relevance of nematodes. Thus, the increased abundance of HSPs in experimental models may reflect conserved cellular mechanisms of response to proteotoxic stress induced by α-synuclein.

In this context, we hypothesize that the significant increase in HSPs abundance observed in the *C. elegans* PD model may result from the accumulation of misfolded α-synuclein and subsequent disruptions in the proper folding of cellular proteins. This process may lead to a proteostress state characteristic of PD.^63^ As demonstrated by van Ham *et al*., the loss-of-function *grk-1* (ok1239) allele selectively attenuates the formation of α-synuclein inclusions while maintaining similar levels of fusion protein expression.^25^ Rather than acting as an unrelated insult, this GRK-1 loss-of-function alters inclusion kinetics, favoring the persistence of soluble conformers that impose continuous demands on the cellular folding and clearance machinery. Crucially, the substantial convergence between these nematode alterations and the clinical PD substantia nigra proteome indicates that this proteostatic activation is primarily driven by conserved, α-synuclein-dependent proteotoxic stress.

When chaperone capacity is exceeded and proper protein refolding fails, cells rely more heavily on primary protein-clearance mechanisms. This shift highlights the ubiquitin-proteasome system (UPS), the principal pathway responsible for regulated protein degradation and a central component of proteostasis. UPS impairment has been strongly implicated in PD, particularly in the context of toxic α-synuclein aggregation.^64^ The system operates through ubiquitin-mediated tagging of substrates for degradation, followed by proteasomal degradation.^65^ The proteasome is composed of α subunits organized into outer rings that perform critical structural and regulatory roles; among them are the PAS-1, 4, 5, and 6 subunits. In the *C. elegans* PD model, we detected upregulation of PAS-1 (*p* < 0.05), 4 (*p* < 0.05), and 5 (*p* < 0.01), whereas PAS-6 (*p* < 0.001) showed downregulation (Fig. 4H-K).

In *C. elegans*, these PAS proteins act as structural elements that regulate substrate entry into the proteasome’s catalytic site.^65^ Therefore, the upregulation of PAS proteins 1, 4, and 5 may represent compensatory cellular responses to insults resulting from the aggregation and deposition of α-synuclein. Furthermore, because the *C. elegans* PD model carries the *grk-1* (ok1239) loss-of-function alongside the α-synuclein transgene, we cannot exclude that GRK-1 itself modulates proteasomal dynamics or sensitizes the degradation machinery. While GRK-1 loss has been shown to reduce inclusion formation without diminishing total α-synuclein levels, it may sustain a persistent population of non-sequestered, soluble misfolded conformers that continuously challenge the proteasome.^25^ Rather than reflecting α-synuclein expression in isolation, these proteasomal alterations likely capture an integrated response to diffuse proteotoxic stress within a genetically sensitized background. In turn, this persistent proteostatic burden compromises mitochondrial homeostasis and engages endoplasmic reticulum stress pathways. Together, these interconnected processes further dopaminergic neuronal degeneration and disease progression.^66, 67^ Similar contributory factors are also implicated in the pathology of other neurodegenerative disorders.^68^

This progressive impairment of proteostasis and the accumulating burden of misfolded proteins directly cascade outward to compromise cellular bioenergetics. Highlighting this downstream, the functional enrichment analysis and metabolic pathways identified substantial proteins from the *C. elegans* PD model associated with BP, and KEGG terms related to the TCA cycle (Fig. 3A-B; Table S4), and CC annotation for cytoplasmic and mitochondrial constituent proteins (Fig. 3A-B; Table S4).

Mitochondrial dysfunction is widely recognized as a key contributor to the pathogenesis of PD. Notably, the core comparative dataset derived from *Jang et al.* PD substantia nigra, revealed that the mitochondrial pathway is the most severely compromised.^36^ In PD, neurons exhibit a high bioenergetic demand that depends on functional mitochondria. Neuronal survival requires continuous adenosine triphosphate (ATP) production to sustain ionic homeostasis, energy-demanding electrical signaling, neurotransmitter release, and vesicular neurotransmitter sequestration.^15^ Furthermore, mitochondria are the primary sources of reactive oxygen species (ROS). When mitochondrial function is compromised, elevated ROS levels accumulate rapidly, driving oxidative stress cascades that ultimately lead to the selective death of dopaminergic neurons in PD.^69^

To assess potential compensatory mechanisms in response to bioenergetic impairment, we analyzed mitochondrial protein regulation. In the *C. elegans* PD model, key metabolic enzymes-including aconitase-2 (ACO-2; *p* < 0.0001) (Fig. 4L), isocitrate dehydrogenase (IDH-1; *p* < 0.001), malate dehydrogenase (MDH-1; *p* < 0.001), acetyl-CoA acyltransferase 2 (ACAA-2; *p* < 0.01) and 3-ketoacyl-CoA thiolase (KAT-1; *p* < 0.01) were upregulated (Fig. 5A-D). Notably, their human orthologs (ACO1, IDH1, MDH1, and ACAT1) were also upregulated in post-mortem substantia nigra tissue from PD individuals. This concordant pattern suggests a conserved metabolic signature across species, indicating a convergence between the transgenic *C. elegans* model and the human disease. However, our analysis also identified additional mitochondrial-associated proteins in the *C. elegans* PD model that were significantly upregulated, including 2-oxoglutarate dehydrogenase (OGDH-1; *p* < 0.01), small ribosomal subunit protein uS9m (MRPS-9; *p* < 0.001), mitochondrial carrier homolog 1 (MTCH-1; *p* < 0.001), and yeast CLU (mitochondrial clustering) related 1 (CLU-1; *p* < 0.01) (Fig. 5E-H). In contrast, their corresponding human orthologs were downregulated.

**Figure 5.**
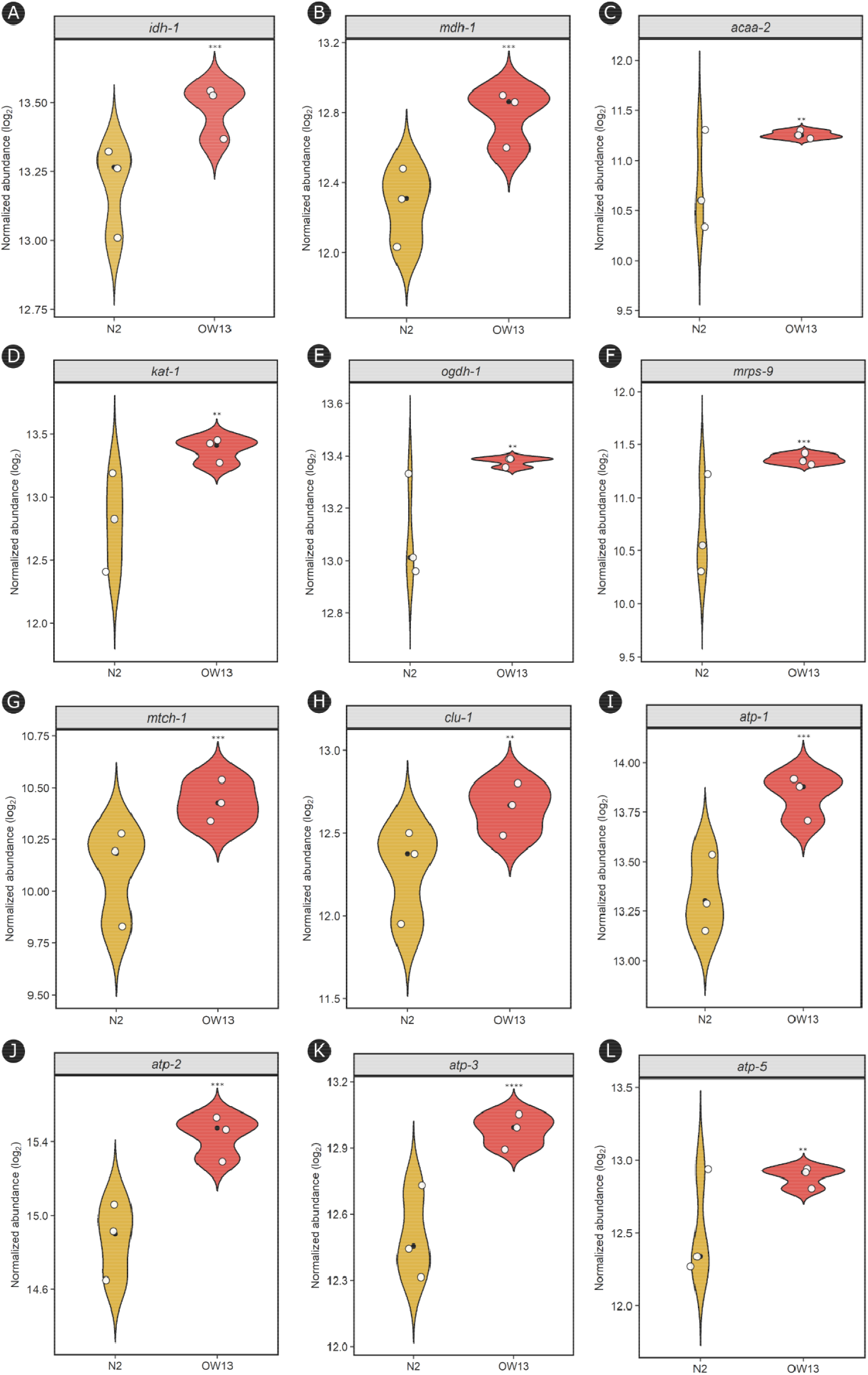
Violin plots of the normalized abundance of DAPs in N2 *wild-type* and *C. elegans* PD model (OW13). (A) Isocitrate dehydrogenase (B) Malate dehydrogenase (C) Acetyl-CoA acyltransferase 2 (D) 3-ketoacyl-CoA thiolase (E) 2-oxoglutarate dehydrogenase (F) Small ribosomal subunit protein uS9m (G) Mitochondrial carrier homolog 1 (H) Yeast mitochondrial clustering related 1 (I) ATP synthase subunit alpha (J) ATP synthase subunit beta (K) ATP synthase subunit (L) ATP synthase subunit 5. Data are displayed as distribution plots, with median values. Statistical significance was assessed using an ANOVA. \*\**p <* 0.01, \*\*\**p <* 0.001, \*\*\*\**p <* 0.0001.

Based on these patterns, we hypothesized that upregulation of mitochondrial enzymes suggests metabolic rewiring towards enhanced activity and energy production. As central components of the TCA cycle and fatty acid β-oxidation, these enzymes indicate a shift in mitochondrial flux, likely acting as a compensatory response to the high energetic and proteostatic demands imposed by α-synuclein aggregation in the *C. elegans* PD model.^70, 71^ This hypothesis provides a molecular framework for secondary phenotypes reported in the literature. For instance, previous studies have reported reduced lipid deposits in the *C. elegans* PD model (OW13) compared with wild-type animals. In this context, the increased abundance of ACAA-2 and KAT-1 raises the hypothesis of potential shifts in fatty acid catabolism, which could provide a plausible molecular rationale for the reduced lipid storage previously described in this PD model.^27, 72^ This metabolic redirection toward lipid utilization reflects a conserved, adaptive response to energetic stress, which is consistent with alterations in lipid metabolism described in PD.^73,74^

At the sub-molecular level, these enzymatic variations alter the organelle’s fundamental driving forces. Mechanistically, ACO-2 catalyzes the isomerization of citrate to D-isocitrate, IDH-1 promotes the oxidative decarboxylation of D-isocitrate to α-ketoglutarate, and MDH-1 catalyzes the oxidation of malate to oxaloacetate, generating NADH that subsequently fuels the electron transport chain.^75–77^ The increased abundance of these proteins in the *C. elegans* PD model suggests the activation of energy-production pathways as a compensatory response to α-synuclein-induced mitochondrial dysfunction. This interpretation aligns with established biochemical frameworks; previous studies have demonstrated that α-synuclein interacts with mitochondria, promoting membrane depolarization and severely reducing ATP production, thereby compromising cellular bioenergetics.^13^

Similarly, the dataset from PD individuals revealed downregulation of mitochondrial-associated proteins, including OGDH, MRPS9, MTCH1, and CLUH, indicating impaired energy metabolism and disrupted mitochondrial redox homeostasis.^36, 78, 77^ In contrast, the corresponding upregulated orthologs in *C. elegans* suggest the activation of compensatory stress-response mechanisms to preserve mitochondrial function, restore cellular bioenergetic balance, and maintain proteostasis under disease-associated conditions.^80^

Consistent with potential bioenergetic remodeling, ATPase levels in the *C. elegans* PD model were also altered, reflecting previously identified mitochondrial changes. In particular, several subunits of mitochondrial ATP synthase (Complex V), including ATP synthase subunit alpha (ATP-1; *p* < 0.001), ATP synthase subunit beta (ATP-2; *p* < 0.001), ATP synthase subunit (ATP-3; *p* < 0.0001), and ATP synthase subunit 5 (ATP-5; *p* < 0.01), were significantly upregulated in the *C. elegans* (Fig. 5I-L). These proteins constitute the core rotational machinery of the oxidative phosphorylation system, which synthesizes ATP using the proton gradient generated by the electron transport chain across the inner mitochondrial membrane.^81^

The increased abundance of ATP synthase subunits complements the observed upregulation of TCA cycle and fatty acid β-oxidation enzymes, suggesting a potential concerted upregulation of mitochondrial pathways involved in reducing-equivalent generation and ATP synthesis in the *C. elegans* PD model. We hypothesize that this pattern may reflect a robust, adaptive attempt to sustain energy homeostasis under α-synuclein-induced mitochondrial stress. The absence of these specific orthologs in the PD individuals dataset may underscore temporal differences between disease models. We propose that while the *C. elegans* PD model captures a compensatory phase, human end-stage samples reflect a moment where these regulatory mechanisms are no longer operative. Another proteomic study in PD supports this compensatory trend, reporting increased abundances of the human orthologs ATP5F1A, ATP5F1B, and ATP5PO in the substantia nigra.^82^ This reinforces our hypothesis that the bioenergetic remodeling observed in *C. elegans* mirrors the molecular changes occurring in the PD brain.

A potential outcome of this hypothesized bioenergetic remodeling and associated mitochondrial stress is an elevated burden of ROS production. Accompanying this internal stress, proteins involved in redox balance showed significant alterations in abundance in our animal model. Glutathione-disulfide reductase (GSR-1; *p* < 0.001) was upregulated in the *C. elegans* PD model (Fig. S3) and PD individuals. This enzyme was associated with the terms cytoplasm and mitochondria in CC (Fig. 3A; Table S4), highlighting its broad involvement in maintaining the integrity of these cellular compartments. Furthermore, superoxide dismutase (SOD-1; *p* < 0.0001) and catalase 1 (CTL-1; *p* < 0.05) were also upregulated in *C. elegans*, pointing to a coordinated increase in the abundance of core antioxidant enzymes (Fig. S3).

These alterations in redox homeostasis are particularly relevant to PD pathophysiology, as oxidative stress is closely associated with synaptic dysfunction and progressive neuronal degeneration.^83^ Mitochondria are known to be important producers of O2·−; however, other sources of ROS, such as the endoplasmic reticulum, result from the action of oxidoreductases, which catalyze the formation of disulfide bridges in proteins.^83^ In this context, the GSR-1, SOD-1, and CTL-1 act in redox regulation in *C. elegans*. GSR-1, the human ortholog of thioredoxin reductase 1 (TXNRD1), is essential for regulating the response to oxidative stress.^84^ The thioredoxin system is an essential antioxidant system; previous studies in PD models have shown that increased TXNRD1 levels reduce endoplasmic reticulum stress and ROS production.^85^ Therefore, we hypothesize that the increased GSR-1 abundance identified in our analysis represents another functional parallel to the *C. elegans* PD model and the human condition, particularly because α-synuclein accumulation is a major contributor to ROS production.

In addition to GSR-1, the coordinated regulation of SOD-1 and CTL-1 highlights the engagement of complementary antioxidant mechanisms to limit oxidative damage. Although we did not detect their human orthologs in the comparative proteomic dataset of PD individuals, these proteins remain important regulators of oxidative stress in *C. elegans*. SOD-1 catalyzes the conversion of superoxide radicals into less toxic species,^86^ and has been associated with increased stress resistance and lifespan in *C. elegans*.^87^ CTL-1 functions as a cytosolic catalase that detoxifies hydrogen peroxide, thereby contributing to the maintenance of cellular redox homeostasis.^88, 89^ The increased abundance of CTL-1 observed in the *C. elegans* PD model is consistent with its established role in the adaptive response to oxidative stress under pathological conditions. Collectively, these findings strengthen the translational relevance of the *C. elegans* model and provide insight into the interplay between protein aggregation and oxidative stress in PD pathophysiology.

The identification of coordinated molecular responses, including alterations in mitochondrial bioenergetics, proteostasis, and the activation of antioxidant defense mechanisms, prompted a broader evaluation of pathway conservation between the *C. elegans* PD model and PD individuals. This analysis sought to determine whether the observed cross-species similarities extend beyond individual orthologous proteins to encompass conserved molecular processes and biological pathways associated with PD pathophysiology. Using an integrated pathway analysis, we demonstrated consistency across species. Thus, the functional terms motor proteins, ribosome, carbon metabolism, TCA cycle, and protein processing in endoplasmic reticulum were enriched in common for proteins from the PD model and PD individuals (Table 1).

**Table 1.** Functional comparison of pathways associated with the *C. elegans* PD model and their corresponding human orthologs identified in PD individuals using a *p* < 0.05 in DAVID.

| Pathways | *C. elegans* ($p$-value) | Protein count* | Human ($p$-value) | Protein count** |
| --- | --- | --- | --- | --- |
| Motor proteins | $p < 0.0001$ | 8 | $p < 0.001$ | 7 |
| Ribosome | $p < 0.0001$ | 11 | $p < 0.01$ | 13 |
| Carbon metabolism | $p < 0.001$ | 8 | $p < 0.0001$ | 7 |
| TCA cycle | $p < 0.01$ | 4 | $p < 0.001$ | 4 |
| Protein processing in endoplasmic reticulum | $p < 0.05$ | 6 | $p < 0.05$ | 5 |

\**C. elegans*; \*\*Human

Among the enriched pathways shared across organisms, the TCA cycle and ER protein-processing pathways stand out. Neurodegenerative diseases, such as PD, can be considered an energy disorder, making neurons vulnerable to oxidative stress and predisposed to neuronal death.^90^ Therefore, based on our comparative findings, the conservation and sharing of the TCA cycle pathway between the *C. elegans* PD model (OW13) and PD individuals are relevant links that connect the model to human disease. Beyond its role in cellular energy production, the TCA cycle is increasingly recognized as a central link between carbon metabolism and redox homeostasis, both of which are critically disrupted during the progression of PD.^91^ Characterizing this shared pathway allows us to observe metabolic similarities between the PD model and PD individuals, supporting its use to study bioenergetic aspects in a mimetic model of energy metabolism disruption under α-synuclein stress.

Beyond bioenergetic processes, the shared enrichment of the ER protein-processing pathway highlights a parallel vulnerability in structural maintenance. The ER plays an important role in protein folding and quality control, and its dysfunction has been associated with the toxic accumulation of misfolded proteins and activation of cellular stress response in neurodegenerative diseases.^92^ From this perspective, the overlapping enrichment of ER protein processing indicates conserved alterations in proteostasis, illustrating that the transgenic nematode experiences a cellular proteostress pattern that mirrors that of PD.^93^

This combined metabolic and proteostatic stress extends beyond the cell body, compromising structural integrity and trafficking networks. Progressive neurodegenerative diseases exhibit disruptions in axonal transport that contribute to disease progression. Motor proteins, including microtubule-associated families, form the vital intracellular transport pathways that enable the trafficking of vesicles, proteins, and organelles.^94, 95^ Dysfunctions in the organization and transport system can compromise intracellular transport, leading to protein accumulation, cell stress, and synaptic communication failures in the nervous system.^96^ Furthermore, evidence indicates that toxic α-synuclein disrupts microtubule dynamics and motor protein function.^97^ In this context, the shared enrichment of motor protein pathways in both the *C. elegans* PD model and PD individuals supports their translational relevance and suggests conservation of cytoskeletal dysfunction across species.

While functional enrichment analysis indicates that the *C. elegans* PD model and PD individuals share perturbed physiological pathways, proteins do not act in isolation but function within coordinated interaction networks. To move beyond static pathway overlap toward a systems-level understanding of molecular connectivity, we therefore analyzed PPI networks constructed from the overlapping cross-species candidates identified in the preceding analysis. In these networks, nodes represent functional proteins and edges represent predicted interactions. Restricting the analysis to the 76 *C. elegans* DAPs and their 70 corresponding human orthologs in PD enabled direct comparison of disease-associated network topology across species (Fig. 6E). Our analyses revealed that the PPI network of the *C. elegans* PD model consists of 76 nodes and 386 edges (Fig. 6A). To evaluate signal propagation in this network, we computed topological properties reflecting its structural organization and connectivity.^98^ The “*hub*” proteins of a network are mathematically defined as the nodes with the highest degree of connectivity and tend to play an essential role in regulating the PI network.

**Figure 6.**
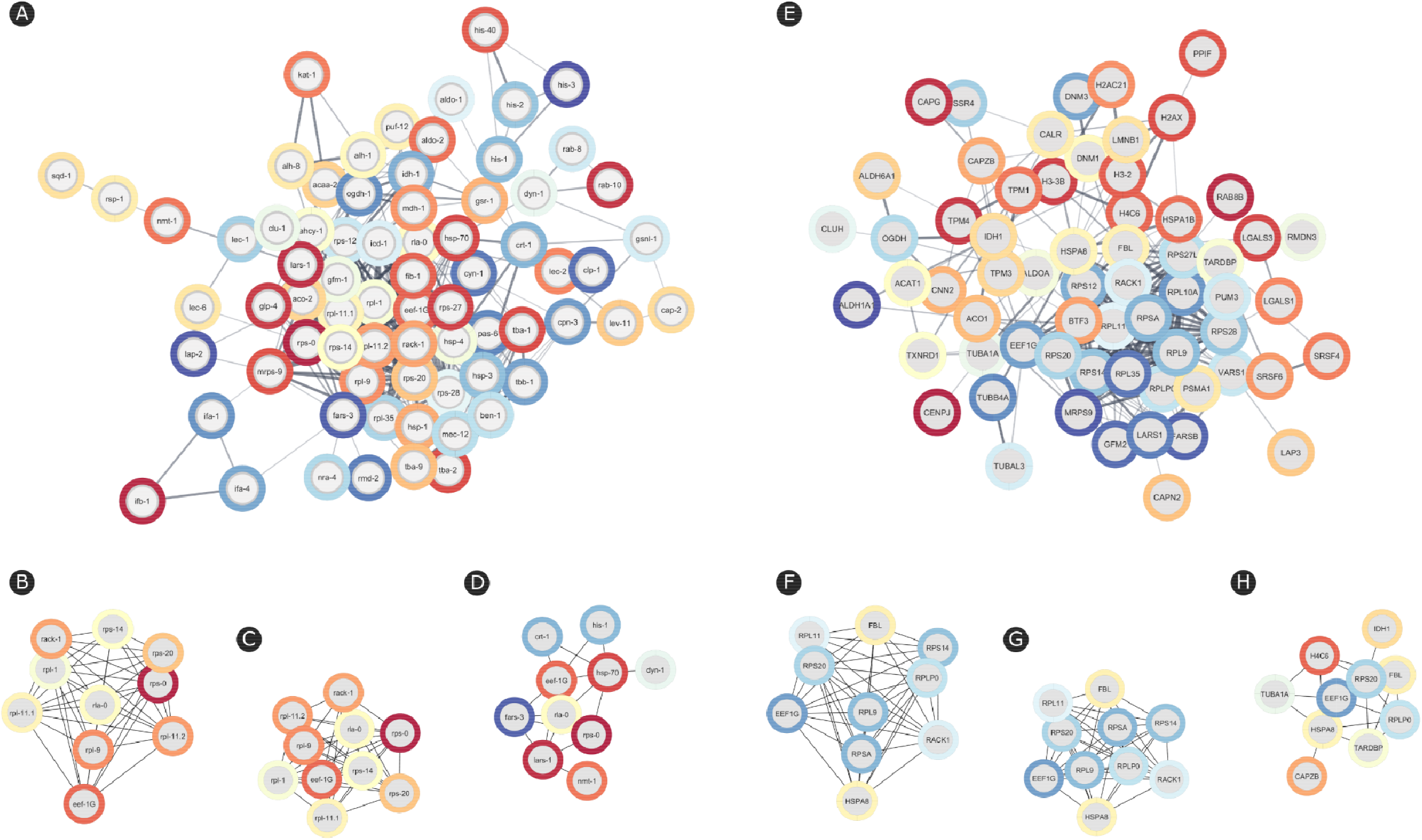
PPI network from the *C. elegans* PD model (OW13) and PD individuals’ proteins of cross-species analysis. (A) PPI network of the *C. elegans* PD model. (B-D) Degree, closeness, and betweenness of the PD model, respectively. (E) PPI network of PD individuals. (F-H) Degree, closeness, and betweenness of PD individuals, respectively. Color scale: Node color gradient denotes expression fold change, where blue shades represent downregulated proteins, yellow/orange indicates intermediate upregulation, and dark red denotes strongly upregulated proteins.

According to Degree, in the PPI network of the *C. elegans* PD model, the proteins EEF-1G, RPS-0, RLA-0, RPL-9, RPS-20, RPL-11.1, RPL-11.2, RPS-14, RACK-1, and RPL-1 were the most relevant hubs (Fig. 6B). The analysis of the topological properties of the C_C_ and C_B_ PPI networks, based on centrality measures, is presented in Figure 6C-D and Table 2.

**Table 2.** Top proteins with the highest degree, C_B_ and C_C_ in the *C. elegans* PD model PPI network.

| S. no. | Name | Degree | Name | C<sub>B</sub> | Name | C<sub>C</sub> |
| --- | --- | --- | --- | --- | --- | --- |
| 1 | **EEF-1G** | 39.0 | **EEF-1G** | 1321.0 | **EEF-1G** | 53.7 |
| 2 | RPS-0 | 33.0 | **RLA-0** | 682.1 | RPS-0 | 49.8 |
| 3 | **RLA-0** | 31.0 | RPS-0 | 599.5 | **RLA-0** | 48.9 |
| 4 | RPL-9 | 27.0 | HSP-70 | 541.5 | RPL-9 | 46.6 |
| 5 | RPS-20 | 25.0 | LARS-1 | 502.7 | RPS-20 | 45.6 |
| 6 | RPL-11.1 | 23.0 | FARS-3 | 441.0 | RPS-14 | 44.6 |
| 7 | RPL-11.2 | 23.0 | HIS-1 | 408.0 | RACK-1 | 44.0 |
| 8 | RPS-14 | 23.0 | CRT-1 | 339.2 | RPL-11.1 | 44.0 |
| 9 | RACK-1 | 23.0 | DYN-1 | 295.4 | RPL-11.2 | 44.0 |
| 10 | RPL-1 | 22.0 | NMT-1 | 276.0 | RPL-1 | 43.8 |

Identification of high-centrality hubs is critical, as they represent key regulators occupying central positions that enable broad influence over downstream functional processes.^99, 100^ To identify the robust network regulators in the *C. elegans* network, we intersected the top-ranked nodes across three independent topological measures (degree, C_C_, and C_B_) using a Venn diagram (Fig. S4). This multi-metric filtering yielded three shared nodes, EEF-1G, RPS-0, and RLA-0, indicating that these proteins constitute a central structural core potentially involved in coordinating the response to α-synuclein stress. Consistent with their centrality, all three proteins were significantly upregulated, with EEF-1G (*p* < 0.01), RPS-0 (*p* < 0.0001), and RLA-0 (*p* < 0.0001) (Fig. S3).

To evaluate whether human PD brain pathology exhibits a similar structural organization, we applied the same mathematical filtering procedure to the PD network, comprising 69 nodes and 299 edges (Fig. 6E). According to Degree, the hub proteins EEF1G, RPS20, HSPA8, RPLP0, RPL9, RPSA, RPL11, RPS14, FBL, and RACK1 were identified in the PPI network (Fig. 6F). Topological properties including C_C_ and C_B_ were also analyzed, identifying 5 key regulators familiar to Degree, C_C_, and C_B_: EEF1G, RPS20, HSPA8, RPLP0, and FBL (Fig. 6G-H, Table 3).

**Table 3.** Top proteins with the highest degree, C_B_ and C_C_, in the PD individuals.

| S. no. | Name | Degree | Name | $C_B$ | Name | $C_C$ |
| --- | --- | --- | --- | --- | --- | --- |
| 1 | **EEF1G** | 26.0 | HSPA8 | 615.6 | RPS20 | 42.8 |
| 2 | RPS20 | 26.0 | **EEF1G** | 450.2 | HSPA8 | 42.6 |
| 3 | HSPA8 | 25.0 | IDH1 | 378.6 | **EEF1G** | 42.0 |
| 4 | **RPLP0** | 25.0 | TARDBP | 278.6 | **RPLP0** | 41.6 |
| 5 | RPL9 | 24.0 | TUBA1A | 249.8 | RPL9 | 41.3 |
| 6 | RPSA | 24.0 | RPS20 | 244.0 | RPSA | 41.2 |
| 7 | RPL11 | 23.0 | FBL | 237.5 | RPL11 | 41.0 |
| 8 | RPS14 | 23.0 | H4C6 | 216.5 | RPS14 | 41.0 |
| 9 | FBL | 22.0 | CAPZB | 192.1 | FBL | 39.5 |
| 10 | RACK1 | 20.0 | **RPLP0** | 187.7 | RACK1 | 39.5 |

The key regulators of PPI networks in the *C. elegans* PD model and in PD individuals were compared to assess overlap in their regulation. The proteins EEF-1G (EEF1G) and RLA-0 (RPLP0), therefore emerged as shared key regulators across the species. The sharing of these key regulators suggests the presence of common molecular mechanisms associated with the regulation of protein networks. Thus, the findings reinforce the relevance of the *C. elegans* PD model for investigating the molecular processes underlying the disease, highlighting a shared regulatory process that brings the experimental model closer to the pathophysiology observed in human brains.

To investigate the local functional organization of these master regulators, we performed a cluster-based module analysis to partition the *C. elegans* network into functional subcomplexes. Modules correspond to groups of proteins that display a higher frequency of interactions among themselves.^45^ The analysis of module dynamics can provide important insights into how biological systems adapt to different conditions. Therefore, the initial native network was divided into modules using the MCODE plugin in Cytoscape 3.10.4, which identified 7 significant modules with MCODE Scores ≥ 2. The key regulatory protein EEF-1G was identified as a component of module 1, the highest-scoring module (Score: 17.7), with 19 nodes and 160 edges (Fig. S5).

In *C. elegans*, EEF-1G constitutes the gamma subunit of the eukaryotic elongation factor 1 (eEF1) complex, responsible for adding amino acids to nascent polypeptide during elongation, and is essential for protein synthesis.^101^ Proteomic analysis of the *C. elegans* PD model revealed upregulation of EEF-1G (*p* < 0.01) and RLA-0 (*p* < 0.0001), whose human orthologs are EEF1G (eEF1Bγ) and RPLP0, respectively. Members of the eEF1 complex are recognized as key modulators of neuronal homeostasis, with alterations in the translation elongation machinery linked to neuronal dysfunction.^102^

In humans, the eEF1Bγ (encoded by the EEF1G gene) appears to play a structural role within the elongation complex, acting as a scaffold that anchors components of the eEF1 machinery to the cytoskeleton and enhancing translational efficiency by facilitating GDP exchange on eEF1Bα.^102^ In contrast to the *C. elegans* PD model, the proteomic dataset from PD individuals revealed decreased abundance of both eEF1Bγ and RPLP0, suggesting a divergent regulation of translational control between the experimental model and the human condition. Notably, reduced levels of eEF1 complex components, including eEF1A, have previously been associated with PD progression, further supporting the involvement of translation dysregulation in disease pathology.^103^

Ultimately, defining this conserved translational and metabolic landscape provides a structured molecular framework to guide future preclinical investigations. Future perspectives for deepening the molecular aspects of the *C. elegans* PD model (OW13) include experimentally evaluating the hypotheses raised by our study that support and strengthen the model’s use in the study of PD, to gain a deeper understanding of still-obscure aspects of its pathophysiology. Furthermore, it is necessary to correlate the findings with potential behavioral phenotypes that enable the phenotypic observation of the molecular manifestations of disturbances generated by the accumulation and aggregation of α-synuclein.

## 4. Limitations

Our study presents some limitations that should be considered when interpreting the findings. Although *C. elegans* provides a valuable platform for investigating conserved molecular mechanisms, it does not fully recapitulate the complexity of the human nervous system involved in PD. In addition, the experimental model is based on α-synuclein expression in body wall muscle cells rather than in dopaminergic neurons, which may limit the direct translation of the results to neuronal pathology. The OW13 strain harbors the *grk-1* (ok1239) loss-of-function alongside the integrated *pkIs2386* transgene. Because GRK-1 functions as an endogenous genetic modifier of α-synuclein inclusion formation, comparing OW13 against N2 animals captures the combined proteomic consequences of α-synuclein proteotoxicity and GRK-1 deficiency, rather than α-synuclein expression alone.

The cross-species comparison also involves distinct biological contexts, as the human data were derived from post-mortem substantia nigra tissue, whereas the *C. elegans* proteome reflects whole-organism analysis, potentially masking tissue-specific effects. Furthermore, the proteomic approach employed is inherently descriptive and does not establish causal relationships, underscoring the need for further functional validation of the proposed mechanisms. Finally, some hypotheses raised in this study, particularly those related to lipid metabolism and bioenergetic remodeling, remain to be experimentally validated, highlighting the need for targeted studies to confirm these observations.

## 5. Conclusion

Our study provides the first comprehensive proteomic characterization of the transgenic *C. elegans* PD model (OW13) expressing α-synuclein and demonstrates the existence of conserved molecular alterations shared with the Parkinsonian substantia nigra. Cross-species comparative analysis identified orthologous proteins and pathways associated with mitochondrial metabolism, proteostasis, oxidative stress regulation, ribosomal function, and endoplasmic reticulum protein processing, reinforcing the model’s translational relevance. The enrichment of pathways related to the tricarboxylic acid cycle, mitochondrial function, and protein quality control suggests that α-synuclein induces a broad proteostatic and bioenergetic remodeling response in *C. elegans* that resembles key aspects of PD pathophysiology. Furthermore, the identification of shared key regulators, including EEF-1G/EEF1G and RLA-0/RPLP0, highlights conserved protein interaction networks that may be involved in disease-associated cellular adaptation.

The findings also support the hypothesis that the molecular alterations observed in the *C. elegans* PD model may reflect an early compensatory response to proteotoxic and mitochondrial stress. The upregulation of heat shock proteins, antioxidant enzymes, TCA cycle components, and ATP synthase subunits indicates activation of protective mechanisms that maintain proteostasis, redox balance, and energy homeostasis under α-synuclein-induced stress. Altogether, these findings underscore the value of *C. elegans* as a complementary experimental system for investigating conserved molecular mechanisms underlying PD and for generating data-driven hypotheses regarding the interconnected cascades of mitochondrial dysfunction, oxidative stress, and proteostasis imbalance.

## Author Contributions

**I.C.B.:** conceptualization, writing, original draft, reviewing, editing, carrying out the experiment, and data analysis. **M.L.L.V., J.B.F., M.G.A.M., R.A.S.:** writing, original draft, reviewing, editing, and carrying out the experiment. **J.L.L.F., G.J.S.P., P.G.:** reviewing, editing, found acquisition, project administration, and supervision.

## Acknowledgments

We thank FACEPE - Fundação de Amparo à Ciência e Tecnologia de Pernambuco (FACEPE, IBGP-0404-2.12/23; APQ-1223-2.05/22; FACEPE-APQ-1601-4.03/25). The National Council for Scientific and Technological Development (CNPq) for financial support of the National Institute of Science and Technology on Molecular Science (INCT-CiMol - 406804/2022-2) and Universal (404023-2021-5; 406820/2023-6). Also, the Coordenação de Aperfeiçoamento de Pessoal de Nível Superior (CAPES, 88887.956451/2024–00). The Keizo Asami Institute (iLIKA) for providing research infrastructure. Some strains were provided by the CGC, which is funded by the NIH Office of Research Infrastructure Programs (P40 OD010440).

## Abbreviations

aa: amino acid(s)
ACAA-2 / ACAT1: acetyl-CoA acyltransferase 2
ACO-2 / ACO1: aconitase-2
ANOVA: analysis of variance
ATP: adenosine triphosphate
BP: biological process
*C. elegans*: *Caenorhabditis elegans*
C_B_: betweenness centrality
C_C_: cellular component / closeness centrality
CGC: *Caenorhabditis* Genetics Center
CHAPS: 3-[(3-cholamidopropyl)dimethylammonio]-1-propanesulfonate
CLU-1 / CLUH: yeast CLU (mitochondrial clustering) related 1
CTL-1: catalase 1
CV: coefficient of variation
DAP / DAPs: differentially abundant protein(s)
DAVID: Database for Annotation, Visualization and Integrated Discovery
DTT: dithiothreitol
*E. coli*: *Escherichia coli*
eEF1 / eEF1Bγ: eukaryotic elongation factor 1
ER: endoplasmic reticulum
FC: fold change
GO: Gene Ontology
GSR-1: glutathione-disulfide reductase
HSP / HSPs: heat shock protein(s)
IDH-1 / IDH1: isocitrate dehydrogenase
KAT-1: 3-ketoacyl-CoA thiolase
KEGG: Kyoto Encyclopedia of Genes and Genomes
MCODE: Molecular Complex Detection
MDH-1 / MDH1: malate dehydrogenase
MF: molecular function
MRPS-9 / MRPS9: small ribosomal subunit protein uS9m
MTCH-1 / MTCH1: mitochondrial carrier homolog 1
NGM: nematode growth medium
nUPLC-MS/MS: nano-ultraperformance liquid chromatography-mass spectrometry/mass spectrometry
OGDH-1 / OGDH: 2-oxoglutarate dehydrogenase
PCA: principal component analysis
PD: Parkinson’s disease
PPI: protein-protein interaction
Q-ToF: quadrupole time-of-flight
ROS: reactive oxygen species
SOD-1: superoxide dismutase
TCA cycle: tricarboxylic acid cycle
TXNRD1: thioredoxin reductase 1
UPLC: ultra-performance liquid chromatography
UPS: ubiquitin-proteasome system
YFP: yellow fluorescent protein

